# Molecular Logic of Cell Diversity and Circuit Connectivity in the REM Sleep Hub

**DOI:** 10.1101/2025.10.01.679602

**Authors:** SK Pintwala, JJ Fraigne, B Dugan, E Arrigoni, JH Peever, PM Fuller

## Abstract

The complexity of the brain arises from the diversity of its circuits and the molecular heterogeneity of the cells that compose them. A mechanistic understanding therefore requires mapping cellular identity and connectivity at single-cell resolution. Here we define the cellular taxonomy of the murine sublaterodorsal tegmental nucleus (SLD), a critical hub for REM sleep, using single-nucleus RNA sequencing. We identified all major brain cell classes, with oligodendrocytes as the most abundant, and resolved seventeen transcriptionally distinct neuronal groups defined by neurotransmitters, neuromodulators, and neuropeptides, each with unique molecular signatures. Projection-specific analysis further revealed that glutamatergic subpopulations targeting the ventrolateral periaqueductal gray (vlPAG) and ventral medulla are molecularly distinct, marked by characteristic receptor motifs. Strikingly, we provide the first direct evidence that SLD^GLUT^ neurons innervate the vlPAG. This newly uncovered SLD^GLUT→vlPAG^ pathway represents a previously unrecognized circuit node for REM sleep regulation, with the potential to act as a REM-OFF population suppressing it. Together, these findings establish a transcriptionally resolved atlas of the SLD, reveal the molecular logic of its circuit connectivity, and nominate candidate molecular actuators of REM sleep control, opening new avenues for dissecting how brainstem circuits orchestrate REM state and its transitions.

## Introduction

Neurons are defined by their anatomical, functional and neurochemical properties, but are most often distinguished by the expression of neurotransmitters, neuromodulators or neuropeptides. Advances in single cell RNA sequencing (scRNA-seq) have revealed remarkable molecular diversity, showing that even neurochemically homogenous populations contain multiple subtypes.^1,2^ While scRNA-seq provides unparalleled insight into transcriptome diversity,^3^ it inherently lacks spatial context. Spatial transcriptomic approaches^4^ overcome this issue by mapping gene expression *in situ* but often overlook the anatomical organization of interconnected neuronal populations.

Neural circuits in the central nervous system (CNS) are organized through long-range axonal projections that form the structural framework through which the CNS mediates distinct neurobiological processes.^5,6^ Within brain regions, neuron types are frequently intermingled but projectionally distinct^7,8^ as individual neurons typically make dedicated projections to specific targets and can be defined by their molecular identity and projection pathways.^9–13^ When taken together, these findings suggest that neuronal organization is intrinsically defined by both transcriptomic and protectional features. Despite advances in profiling CNS gene expression at cellular resolution, how transcriptomic identify aligns with projection-defined connectivity remains largely unknown.

Bridging the transcriptome and projectome requires approaches that resolve gene expression in the context of neuronal projections. Projection-based specialization may emerge through differential gene expression, with transcriptional programs shaping neuronal morphology,^14,15^ connectivity,^16,17^ and function.^18^ Identifying projection-specific molecular signatures would not only expand our knowledge of cellular composition but also illuminate how the CNS encodes functional architecture through its long-range circuits.

The sublaterodorsal tegmental nucleus (SLD) of the pons is a key regulator of rapid eye movement (REM) sleep and its defining features,^19–23^ strongly active during this behavioral state.^24^ The induction of REM sleep muscle atonia by SLD glutamatergic neurons (SLD^GLUT^)^23^ is thought to be mediated to large extent by descending projections to the ventral medulla (vM, SLD^GLUT→vM^),^22^ a region critical to the generation of REM sleep motor atonia.^25,26^ The substrates of ascending SLD^GLUT^ projections to initiate REM include the caudal laterodorsal tegmental nucleus^22^ but may also include the ventrolateral periaqueductal grey (vlPAG), a region associated with REM sleep supression.^27^ While circuit-level studies have been central to neuroscience,^5,6^ how projection-defined subpopulations within the SLD relate to molecular diversity and ultimately function remains unknown.

Here, we combine single cell RNA sequencing with projection-specific labeling to define the molecular landscape of the SLD. We hypothesize that the SLD is a transcriptionally diverse hub in which vlPAG- and vM-projecting glutamatergic neurons are defined by distinct molecular signatures that encode specialized roles in REM sleep regulation. Together, this approach aims to provide a transcriptional atlas of the murine SLD as well as a molecular code for the specialized roles of SLD neurons in orchestrating REM sleep regulation.

## Results

### SLD cell type composition

To assess the molecular landscape of the murine SLD, we used single nucleus RNA sequencing (snRNA-seq). Tissue was extracted between 12:00 and 14:00, a period of high sleep pressure (light-dark 12:12, ZT0 = 7am).^28^ Using a brain matrix, mouse brains were sectioned between −5.0mm and −6.0mm from bregma, the SLD was isolated using a 0.75 mm tissue biopsy punch and samples were flash frozen on dry ice (**Fig. 1A**). Nuclei isolation and library preparation were performed by 10X Genomics using the Chromium v4 droplet-based assay on the NovaSeq 500 platform (**Fig. 1B**). Across 29,205 nuclei we obtained a mean of 40,931 reads and a median of 3,408 genes per nucleus. After filtration and quality control in Seurat^29,30^ (nFeatureRNA 500-6,500; nCountRNA 500-30,000; %-mitochondrialRNA <3%), 27,659 high-quality nuclei underwent further analysis.

**Figure 1.**
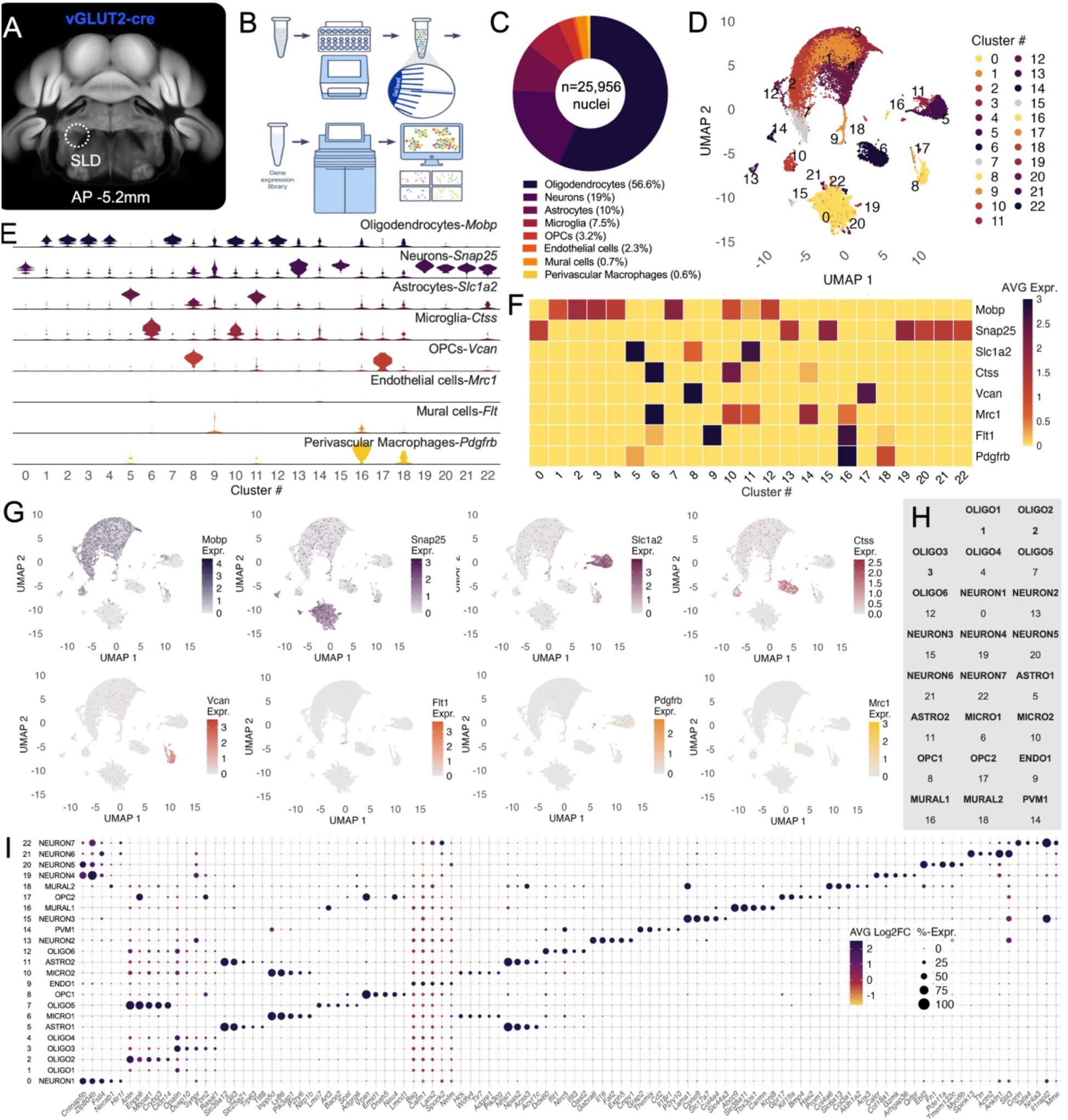
Cell type composition of the murine SLD. **A, B.** The SLD was microdissected from *vGLUT2*-cre mice (n=8, male) and processed using 10X Genomics for snRNA-seq. **C**. Eight cell types were detected in the SLD (n=27,659 nuclei), with oligodendrocytes being the most abundant. **D**. Unsupervised clustering identified 23 transcriptional clusters. **E, F**. Distribution and expression levels of cell-type markers across transcriptional clusters. **G**. Feature plots showing expression of cell type markers across the UMAP space. **H**. Dot plot depicting the top five genes expressed per cluster (Log2FC ≥ 0.25; minimum percent 25%; Wilcoxon rank sum test with the Bonferroni correction *p < 0.05).

We first assessed general cell type composition in the SLD using canonical CNS marker genes (**Fig. 1C**). Oligodendrocytes (*Mobp*, 56.6%) were the most abundant, followed by neurons (*Snap25*, 19.0%), astrocytes (*Slc1a2*, 10.0%), microglia (*Ctss*, 7.5%), oligodendrocyte precursor cells (OPCs, *Vcan*, 3.2%), endothelial cells (*Flt1*, 2.3%), mural cells (*Pdgfb*, 0.7%) and perivascular macrophages (PVMs, *Mrc1*, 0.6%).

Principle component analysis (PCA) and unsupervised clustering using the Louvain method identified 23 distinct clusters (**Fig. 1D**). Uniform Manifold Approximation and Projection (UMAP) analysis for dimensionality reduction and visualization revealed segregation into broad but distinct clusters, which we used to explore higher-order cell type organization.

Cell type identities were assigned based on the distribution (**Fig. 1E**) and expression levels (**Fig. 1F**) of canonical cell type marker genes. Distribution was used for initial cluster annotation, with expression levels serving as a guide (e.g., cluster 14). This resulted in six oligodendrocyte clusters (#1, 2, 3, 4, 7, 12), seven neuron clusters (#0, 13, 15, 19, 20, 21, 22), two astrocyte clusters (#5, 11), two microglia clusters (#6, 10), two OPCs clusters (#8, 17), one endothelial cluster (#9), two mural clusters (#16, 18) and one PVM cluster (#14). Feature plots of marker gene expression over the UMAP further validated these annotations (**Fig. 1G**). Clusters were subsequently renamed according to cell type identity: OLIGO1–6, NEURON1–7, ASTRO1–2, MICRO1–2, OPC1–2, ENDO1, MURAL1–2, and PVM1 (**Fig. 1H**).

To further resolve transcriptional identities, we identified the top five marker genes for each cluster (log2 fold-change, log2FC, ≥ 0.25, minimum percent, 25%, Wilcoxon rank-sum test with the Bonferroni correction). OLIGO clusters expressed canonical oligodendrocyte genes such as *Enpp6* and *Opalin*, with unique gene combinations defining each cluster (**Fig. 1I**). NEURON2 nuclei were enriched for *Gabra6*, NEURON3 for *Slc17a7* (vGLUT3) and NEURON4 for *Calcr* and *Qrfpr*, a hypothalamic peptide receptor associated with wakefulness and motor activity that colocalizes with orexin.^31^ ASTRO clusters expressed *Slc39a12* and/or *Gli3*, while MICRO clusters were enriched for *Ly86*. All other clusters similarly expressed canonical markers alongside novel transcripts, further supporting cluster annotations.

Together, our data indicate that all principal cell types of the brain are present in the SLD, with oligodendrocytes predominating. Our findings also provide a detailed view of the SLD’s cellular composition, genetic signatures, and molecular subclassifications.

### SLD neuronal populations

Next, we focused on neurons in the SLD to assess the relative abundance of neuronal subtypes. We selected nuclei expressing the neuronal marker gene *Snap25* and clusters enriched in non-neuronal marker were removed, yielding 4,654 high-quality neuronal nuclei for downstream analysis (**SFig. 1A, B**; see Methods).

We first examined the expression of neurochemical marker genes relevant to the dorsal pons^32^ (**Fig. 2A**). Neuronal nuclei expressing cholinergic (*Chat, Slc5a7*), monoaminergic (*Th, Slc18a2, Slc6a2*), glutamatergic (*Slc17a6, Slc17a7*), GABAergic (*Slc32a1, Gad1, Gad2*), glycinergic (*Slc6a2*) and peptidergic (*Penk, Gal, Pvalb, Sst*) markers were detected, with excitatory (*Slc17a6+*) nuclei being the most abundant (58.0%).

**Figure 2.**
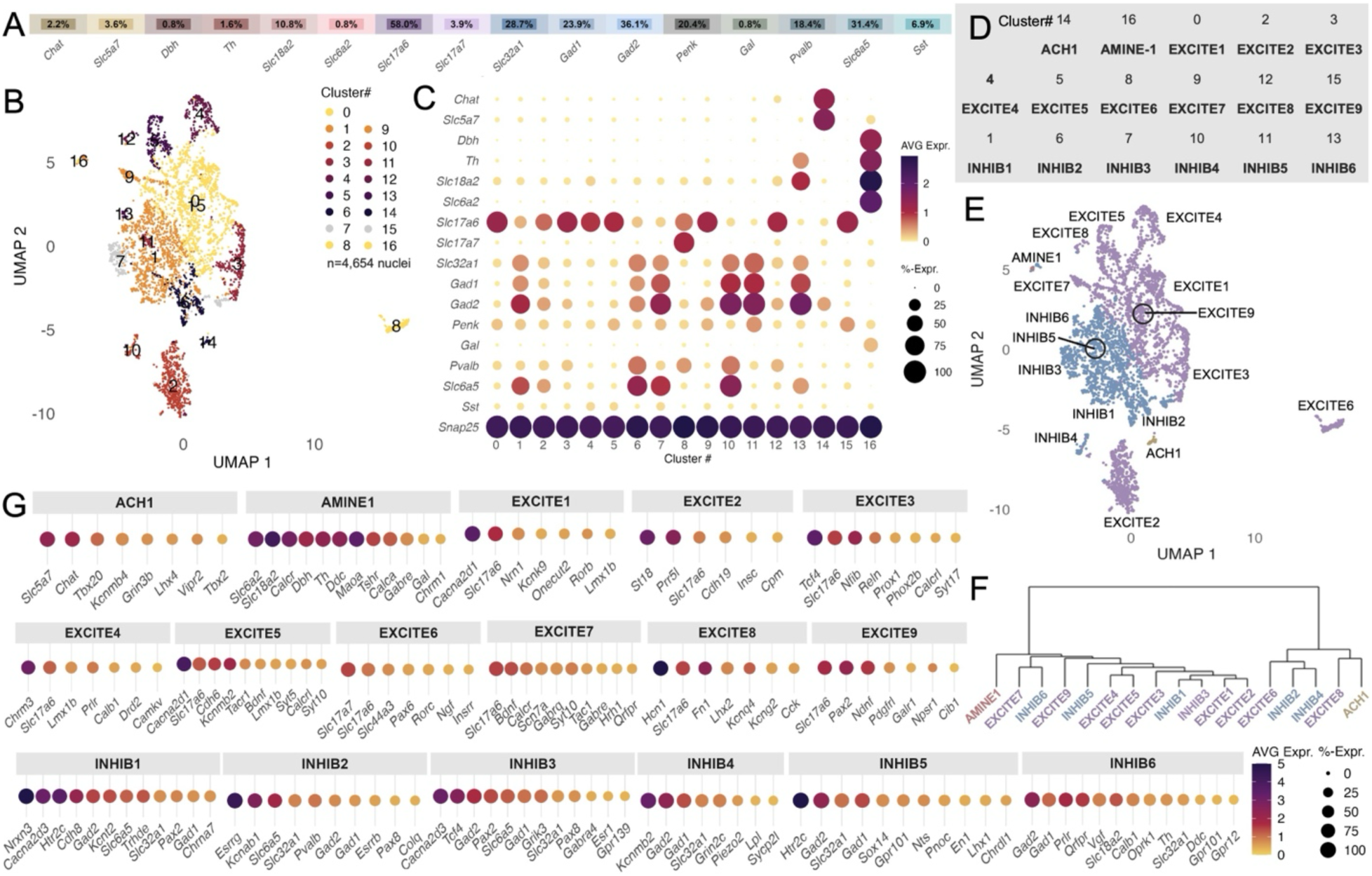
Molecular subtypes of SLD neurons. Neuronal nuclei (n=4,654) were subsetted from the dataset for downstream analysis. **A**. Relative (%-expression) of neurochemical marker genes in all subsetted neuronal nuclei. **B**. UMAP plot of neuronal nuclei forming 17 clusters. **C**. Expression of neurochemical marker genes across neuronal clusters. **D**. Cluster annotations for neurochemical identity. **E**. UMAP plot with neurochemical cluster annotations. **F**. Dendrogram depicting hierarchical organization of neuronal clusters. **G**. Marker genes for each neuronal cluster (Log2FC ≥ 0.25; minimum percent 25%; Wilcoxon rank sum test with the Bonferroni correction *p < 0.05).

Unsupervised clustering with PCA identified 17 distinct neuronal clusters (**Fig. 2B**). Based on neurochemical marker expression (**Fig. 2C**) these were classified as one cholinergic, one monoaminergic, nine glutamatergic and six GABAergic clusters. Parvalbumin and glycine markers largely co-localized with inhibitory (GABAergic) clusters (#1, 6, 7, 10, 13), whereas *Penk* was distributed across both excitatory and inhibitory clusters (#11, 15), indicating overlapping neurochemical expression in SLD neurons. Clusters were annotated as ACH1 (cholinergic) and AMINE1 (monoaminergic), EXCITE1-9 (glutamatergic) and INHIB1-6 (GABAergic) (**Fig. 2D**).

Given that neurochemically-defined clusters were positioned in close proximity on the UMAP (**Fig. 2E**) we performed a hierarchical clustering analysis which revealed that transcriptional profiles did not segregate by neurochemical identity; rather clusters expressing different neurotransmitters and neuromodulators were intermingled (see dendrograms, **Fig. 2F**). This suggests that SLD neurons may exhibit convergent transcriptional profiles shaped by developmental trajectories, spatial organization, projection connectivity, or function rather than neurochemical identity alone.

Differential gene expression analysis between clusters (Log2FC ≥ 0.25; minimum percent 25%; Wilcoxon rank sum test with the Bonferroni correction) identified cluster-defining marker genes encoding neuropeptides, neurotransmitter-related enzymes, transporters, neurotransmitter receptors, neuromodulator receptors, G protein-coupled receptors, ion channels, adhesion proteins, and transcription factors (**Fig. 2G**). When manually filtered, we found that ACH1 nuclei expressed cholinergic machinery (*Slc5a7, Chat*) and transcription factors *Tbx20, Lhx4* and *Tbx2*, implicating these molecular targets in cholinergic neuron development. AMINE1 nuclei were enriched in monoaminergic markers (*Slc6a2, Slc18a2, Th)* and additional transcripts including *Calcr, Dbh, Ddc, Maoa, Tshr, Calca, Gabre, Gal*, and *Chrm1*, highlighting a diverse monoaminergic population.

Among excitatory clusters, EXCITE3 expressed *Tcf4, Calcrl* and *Phox2b*, marking a pontine population controlling respiration.^33^ EXCITE4 expressed C*hrm3, Lmx1b, Prlr, Calb1, Camkv*, and *Drd2*, consistent with pontine dopamine-mediated inhibition of REM sleep and its features.^34^ EXCITE6 predominately expressed *Slc17a7* (vGLUT3), *Pax6* and *Ngf*, suggesting multiple glutamate-release mechanisms. EXCITE7 expressed *Bdnf, Calcr, Gabrq, Tac1, Gabre, Hrh1*, and *Qrfpr*. EXCITE8 nuclei expressed *Lhx2* and *Cck*, and EXCITE9 expressed *Pax2* and *Galr1*.

Inhibitory clusters exhibited similarly diverse profiles: INHIB1 expressed *Nrxn3, Htr2c, Slc6a5, Pax2*, and *Chrna7*; INHIB2 expressed *Esrrg* and *Esrrb, Slc6a5, Pax8* and *Pvalb*, potentially identifying a population of SLD interneurons.^35,36^ INHIB3 nuclei expressed *Cacna2d3, Tcf4*, Pax2, *Slc6a5, Pax8, Gabra4* and *Esr1*. INHIB5 expressed *Htr2c, Nts, Pnoc*, and *Lhx1*, while INHIB6 expressed *Prlr, Qrfpr, Oprk1, Slc18a2, Th*, and *Ddc*, identifying a GABAergic population capable of dopamine co-release.

Overall, SLD neurons are neurochemically diverse, expressing overlapping combinations of neurotransmitters, neuromodulators, and neuropeptides. These findings provide a detailed transcriptional map of SLD neuronal populations and reveal novel gene targets for studying the molecular basis of pontine neurobiology.

### Gene expression profiles of projection-specific SLD^GLUT^ neurons

We next examined the gene expression profile of projection-defined subpopulations of SLD^GLUT^ neurons by combining retrograde tracing with snRNA-seq. In *vGLUT2-Cre* mice, cre-dependent retrograde tracers were unilaterally delivered to the vlPAG (AAV2retro-EF1a-DIO-hChR2(H134R)-mCherry) or vM (AAV2retro-CAG-FLEX-rc[Jaws-KGC-GFP-ER2]; **Fig. 3A**; n=8), labelling SLD^GLUT→vlPAG^ and SLD^GLUT→vM^ neurons with mCherry and GFP, respectively. To reliably identify reporter-positive neurons, we generated a custom reference transcriptome containing *mCherry* and *Gfp* transcripts. Glutamatergic neurons were subsetted by *Slc17a6* expression, and clusters enriched in non-neuronal markers were removed (**SFig. 1C, D**), yielding high-quality glutamatergic nuclei for analysis.

**Figure 3.**
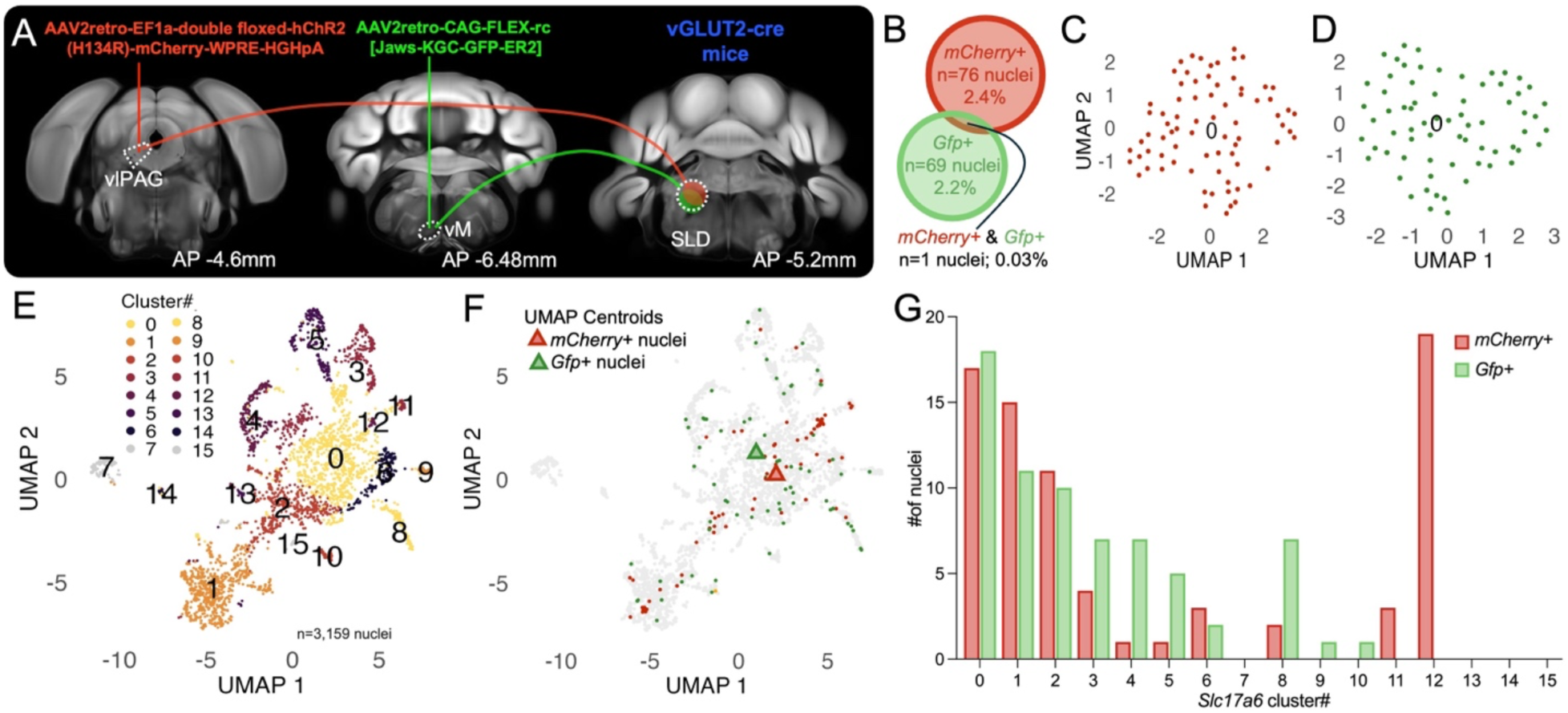
Gene expression profile of vlPAG- and vM-projecting SLD^GLUT^ neurons. **A.** Projection-specific glutamatergic neurons in the SLD were labelled by delivering cre-dependent retrograde tracers to the vlPAG (AAV2retro-EF1a-double floxed-hChR2(H134R)-mCherry-WPRE-HGHpA) and vM (AAV2retro-CAG-FLEX-rc[Jaws-KGC-GFP-ER2]) of vGLUT2-cre mice (n=8). **B**. Proportions of *mCherry+* and *Gfp*+ neuronal nuclei. **C, D**. UMAP plots of *mCherry+* (red) and *Gfp+* (green) neuronal nuclei. **E**. UMAP plot of subsetted *Slc17a6*+ neuronal nuclei (n=3,159). **F**. Feature plot with *mCherry* and *Gfp* expression overlaid. UMAP centroids (triangles) represent the central position of *mCherry+* or *Gfp+* neuronal nuclei on the UMAP plot. **G**. Bar plot with distribution of *mCherry+* or *Gfp+* neuronal nuclei per *Slc17a6* transcriptional cluster.

Of 3,159 *Slc17a6+* neuronal nuclei, 76 expressed *mCherry* (2.4%), 69 expressed *Gfp* (2.2%) and one expressed both (0.03%) (**Fig. 3B**). Thus, vlPAG and vM-projecting SLD^GLUT^ neurons represent small, largely non-overlapping subsets. PCA and unsupervised clustering of *mCherry+* or *Gfp+* nuclei formed single clusters on their respective UMAPs (**Fig. 3C, D**), indicating relative molecular homogeneity within each projection-specific population. PCA and unsupervised clustering of all *Slc17a6*+ nuclei (n=3,179) identified 16 clusters (**Fig. 3E**). Overlaying reporter expression revealed that the labeled nuclei were distributed across multiple clusters but showed distinct UMAP centroid positions (**Fig. 3F**), suggesting transcriptional polarization of projection-defined SLD^GLUT^ neurons. Further supporting this finding, *mCherry+* and *Gfp+* neuronal nuclei were present in different *Slc17a6* transcriptional clusters at varied proportions (**Fig. 3G**). Together, these results suggest that vlPAG and vM-projecting SLD^GLUT^ neurons are transcriptionally intermingled yet polarized, representing divergent states of a common neuron type rather than discrete molecular subtypes.

We next identified differentially expressed genes (DEGs; Log2FC ≥ 0.25; minimum percent 25%; Wilcoxon rank sum test with the Bonferroni correction, *p < 0.05) in projection-specific populations. Compared to *Slc17a6*+ nuclei lacking *mCherry, mCherry*+ nuclei exhibited a total of 673 DEGs (**Fig. 4A**), including genes encoding neurotransmitter receptors, neuromodulator receptors, G protein-coupled receptors, ion channels, adhesion proteins, and transcription factors (**Fig. 4B**). Notably, *Adra1a*, (alpha 1 adrenergic receptor subunit), *Hcrtr2* (hypocretin receptor 2), and *Tacr1* (neurokinin 1 receptor) were enriched in *mCherry*+ nuclei (**Fig. 4B**). In *Gfp*+ nuclei, 1,436 DEGs were detected relative to *Slc17a6+* nuclei lacking *Gfp* (**Fig. 4C**), with enriched transcripts including *Cnr1* (cannabinoid receptor 1), *Glra1* (glycine receptor alpha 1), and *Oprm1* (mu opioid receptor 1, **Fig. 4D**).

**Figure 4.**
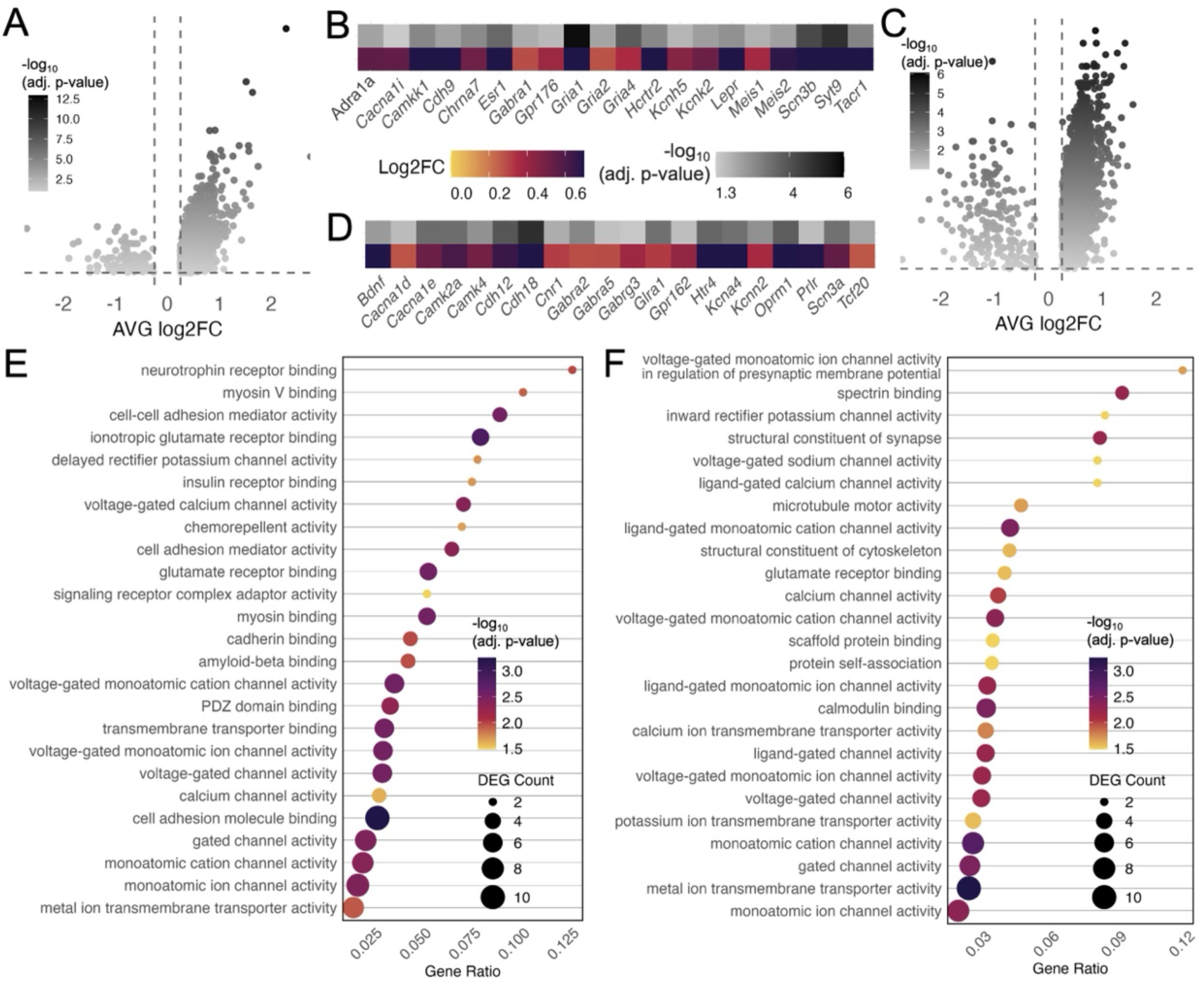
Molecular features of vlPAG- and vM-projecting SLD^GLUT^ neurons. **A.** Volcano plot depicting DEGs for *mCherry+* neuronal nuclei (relative to *Slc17a6+ mCherry-* neuronal nuclei; adjusted p-value, adj. p-value). **B**. Heatmap with 20 select DEGs for *mCherry+* neuronal nuclei. **C**. Volcano plot depicting DEGs for *Gfp+* neuronal nuclei (relative to *Slc17a6+ Gfp-* neuronal nuclei). **D**. Heatmap with 20 select DEGs for *Gfp+* neuronal nuclei. **E, F**. EnrichR data for DEG ontology in *mCherry+* (left) and *Gfp+* (right) nuclei using Molecular Function (MF) categories.

Gene set enrichment analysis (EnrichR) of DEGs using the Molecular Function ontology revealed distinct pathways signatures. *mCherr*y+ nuclei were enriched for categories including neurotrophin receptor binding, cell-cell adhesion mediator activity, ionotropic glutamate receptor binding, voltage gated calcium channel activity and glutamate receptor binding activity (**Fig. 4E**). *Gfp*+ nuclei were enriched for transcripts related to voltage-gated monoatomic ion channel activity in regulation of presynaptic membrane potential, spectrin binding, structural constituents of synapse, voltage-gated sodium channel activity and ligand-gated calcium channels (**Fig. 4F**).

Overall, projection-defined SLD^GLUT^ neurons exhibit graded transcriptional differences along a continuum, rather than forming distinct molecular subtypes. Importantly, projection-specific populations show selective enrichment for genes encoding neurochemical receptors, such as *Hcrtr2* in vlPAG-projecting neurons and *Oprm1* in vM-projecting neurons, suggesting that projection-defined neural populations may be tuned by gene expression programs to receive differential presynaptic inputs. These findings provide insights into how projection-specific SLD^GLUT^ neurons achieve functional specialization in circuit-level processes and behavior.

## Discussion

We present the first single-nucleus transcriptomic atlas of the SLD, a pontine hub essential for REM sleep regulation.^19–23^ Our analyses reveal the major cellular constituents of the SLD, uncover transcriptional diversity within its neuronal population, and define the molecular programs of projection-specific glutamatergic neurons. Taken together, our work develops a conceptual framework uniting transcriptomic and projection identity, further establishing a molecular framework that links gene expression with circuit-level function in REM sleep.

The SLD contains all major CNS cell classes (**Fig. 1C**), with oligodendrocytes as the most abundant, aligning^37^ with previous studies and diverging^32^ from others. Neurons segregated into excitatory, inhibitory, cholinergic, and monoaminergic clusters (**Fig. 2C**), but these clusters did not align neatly with classical neurotransmitter phenotypes (**Fig. 2F**). Instead, transcriptional programs appeared to reflect additional organizing principles such as developmental history,^38,39^ spatial organization,^40^ projection targets^41^— consistent with observations in other brain regions^9,17^ and species.^42^

Within glutamatergic neurons, the principal efferents of the SLD, we show that vlPAG- and vM-projecting subpopulations exhibit partially overlapping but polarized transcriptional profiles. Both subgroups were intermingled in UMAP space (**Fig. 3F**) and distributed across several clusters (**Fig. 3G**), consistent with divergent states of a shared lineage rather than distinct molecular subtypes. Notably, vlPAG-projecting neurons were enriched for *Adra1a, Hcrtr2, and Tacr1* (**Fig. 4B**).

The human sleep disorder narcolepsy is characterized by diminished levels of hypothalamic orexin,^31,43–45^ leading to REM sleep dysregulation, including short latencies to REM sleep.^46^ Our data suggest a mechanism whereby orexin activation of SLD^GLUT→^vlPAG neurons via orexin receptor 2 (Hcrt2r), could drive feed-forward activation of REM-OFF vlPAG GABAergic neurons, a circuit that is weakened with orexin loss. This interpretation aligns with the observation that Hcrt2r in the vlPAG region of an orexin-2-receptor-deficient narcolepsy mouse model suppresses cataplexy while not improving maintenance of wakefulness.^47^

Importantly, the current study provides, to our knowledge, the first direct evidence that SLD^GLUT^ neurons innervate the vlPAG. We consider the possibility that SLD^GLUT→vlPAG^ neurons might constitute a novel REM-OFF population (**Fig. 5**), contributing to pontine REM sleep suppression in conjunction with vlPAG GABAergic projections to the SLD and LC.^27^ However, based on their excitatory nature, their ascending projections to the REM-OFF vlPAG region, and their expression of excitatory orexin receptor 2 (*Hcrtr2*) and alpha 1 noradrenergic receptors (*Adra1a*), we favor the alternative that SLD^GLUT→vlPAG^ neurons are REM-OFF (**Fig. 5**), In this model orexinergic and adrenergic signaling during wakefulness activates SLD^GLUT→vlPAG^ neurons, which in turn drive wake-active, REM-suppressing REM-OFF vlPAG GABAergic neurons that feed back to inhibit SLD REM-ON neurons, thereby preventing REM entry and loss of muscle atonia (i.e. cataplexy). This framework is consistent with evidence that a small population of glutamatergic neurons in this region is maximally active during wakefulness.^24^ and it helps reconcile orexin inputs to the SLD with existing models of pontine REM-sleep control (for review see ^48^). Future experiments will be required to define the precise state dependence and causal role of these SLD^GLUT→vlPAG^ neurons in behavioral-state control.

**Figure 5.**
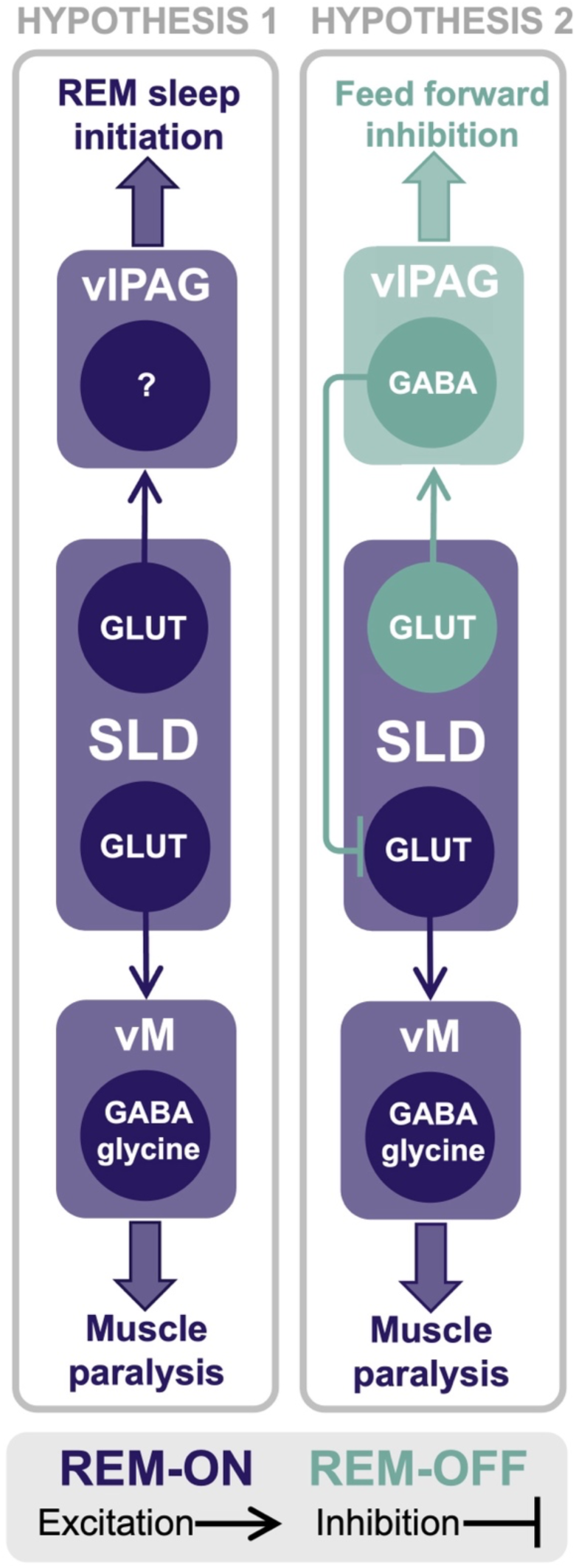
Hypothesized circuit mechanisms of SLD^GLUT^ neurons. In **Hypothesis 1**, both SLD^GLUT→vlPAG^ and SLD^GLUT→vM^ neurons are REM-ON, SLD^GLUT→vlPAG^ neurons engage an unidentified vlPAG population, potentially local inhibitory interneurons, to promote REM sleep, while SLD^GLUT→vM^ neurons activate GABAergic and glycinergic vM neurons to induce muscle paralysis. Together, these circuits drive REM sleep and associated muscle paralysis, respectively. In **Hypothesis 2**, SLD^GLUT→vlPAG^ neurons are REM-OFF and wake active. In this model the recruit vlPAG^GABA^ to provide feed-forward inhibition to SLD^GLUT→vM^ neurons, thereby suppressing REM-related atonia during wakefulness.

Temporally silencing orexin neurons does not induce cataplexy, the pathognomonic symptom of narcolepsy,^49^ suggesting downstream effector(s) redundancy that maintains some level of appropriate regulation in the absence of orexin input. Activation of glutamatergic neurons in the SLD is sufficient to induce cataplexy,^49^ where divergent pathways^22^ are hypothesized to underlie the dissociation of cortical activity and muscle tone^48,50^ through inappropriate activation of SLD^GLUT^ projections to the vM. Our work further identified that vM-projecting SLD^GLUT^ neurons were enriched for modulatory receptors including *Cnr1, Glra1* and *Oprm1* (**Fig. 4D**). Interestingly, opiates increase the number of orexin neurons in the LH,^51^ and suppress cataplexy in both mouse models of narcolepsy^51^ and in human narcolepsy.^52^ Our results identify a mechanism of opioid-mediated suppression of SLD^GLUT→vM^ neurons through *Oprm1* signaling, which may have relevance for cataplexy in narcolepsy. Taken together, the projection-specific receptor repertoires of vlPAG- and vM-projecting SLD^GLUT^ neurons suggest that long-range inputs may differentially tune ascending and descending SLD outputs to regulate REM sleep initiation and motor atonia, respectively.

Taken together, our results indicate that SLD molecular diversity is organized along both cell-type and projectional dimensions. Projection-specific glutamatergic neurons occupy a transcriptional continuum, derived of a common cell type, with selective receptor expression conferring input–output specializations that may underlie the coordination of REM sleep initiation and paralysis. This molecular atlas thus provides a foundation for future studies integrating spatial transcriptomics and physiology to determine how projection-defined transcriptional programs shape SLD activity and REM sleep behavior. By linking molecular profiles to known projection pathways, our results also extend the view of the SLD beyond its anatomical boundaries, suggesting that its contributions to cortical activation and motor regulation arise from a coordinated but diverse cellular repertoire. Elucidating the sources of distinct neurochemical-specific inputs will be essential to understanding the organizational logic of SLD circuitry and how the SLD integrates multiple modes of afferent regulation. Additionally, investigating the transcriptional identity of projection-defined SLD GABAergic^21,22^ neurons would further elucidate the molecular motifs of pontine-driven sleep behavior.

Beyond the SLD, our study also highlights broader methodological considerations. Single-nucleus RNA-seq offers a powerful lens into cellular diversity, but the identification of discrete clusters relies on heuristic assumptions about cell-type boundaries.^1^ Accordingly, we favor the term “transcriptional variants” over “cell types,”^53^ recognizing that gene expression states may represent both dynamic adaptations and stable identities. In this view, our observations regarding projection-based neural populations are *a priori*, based on the foundation of knowledge that neural projections are functional and mediate distinct neurobiological processes.^5,6^ Technical variables—including regional differences in marker gene expression^37^ constraints of barcode-based projectome approaches,^11,54^ and variability in cell recovery^55,56^ further complicate classification. These caveats underscore the need to interpret transcriptomic data within anatomical and functional context rather than in isolation.

Together, these perspectives argue for an integrative framework in which transcriptional states serve as proxies for both intrinsic cellular identity and the selective pressures imposed by circuit connectivity. For the SLD, this approach provides a foundation for linking molecular diversity to its role in sleep–wake regulation. More broadly, it emphasizes the value of combining transcriptomic, anatomical, and projection-based approaches to achieve a deeper understanding of how molecular programs shape neural circuit function.

## METHODS

### Animals

In this study male *vGLUT2-Cre* mice (Slc17a6tm2(cre)Lowl/J on C57BL/6 background, Jackson Laboratory) aged 8-14 weeks were used and bred in-house. Genotyping was performed with polymerase chain reaction using the following primers for Cre-recombinase: forward CACGACCAAGTGACAGCAAT and reverse: AGAGACGGAAATCCATCGCT. All experimental protocols were approved by the University of Toronto’s Animal Care Committee, in accordance with the Canadian Council on Animal Care. Animals were housed under a controlled 12/12 light-dark cycle with food and water provided *ad libitum*.

### Virus preparation for retrograde tracing

For retrograde tracing cre-inducible recombinant adeno-associated viruses (AAVs) were purchased from and packaged by Addgene. One vector carried the transgene mCherry (AAV2retro-EF1a-DIO-hChR2(H134R)-mCherry-WPRE-HGHpA, plasmid #20297, 1 x10^13^ GC/mL) and the other GFP (AAV2retro-CAG-FLEX-[KGC-Jaws-GFP]-ER2; plasmid #84445, 1×10^13^ GC/mL). These vectors employ the AAV2retro helper virus,^57^ which enables efficient retrograde transduction.

### Surgery

Mice were anesthetized using isoflurane (3% induction, 1-2% maintenance in 0.8L/minute of oxygen) until loss of standard reflexes. Meloxicam was administered pre-operatively (20mg/kg, subcutaneous) for post-operative analgesia. For virus infusion surgeries mice were affixed to the stereotaxic frame (model 902; Kopf Instruments). Using a 28-guage cannula connected to a Pump 11 Elite digital microinjection pump (Harvard Apparatus) and p20 tubing, viruses were unilaterally delivered (200nL) to the vlPAG (anterior-posterior: 4.5, medial-lateral: +0.5, dorsal-ventral: −2.6, in mm) and vM (anterior-posterior: −5.0, medial-lateral: +0.9, dorsal-ventral −4.25, in mm) relative to bregma or the pial surface. Hence, glutamatergic SLD neurons projecting to the vlPAG and vM were retrogradely labelling via mCherry and GFP, respectively.

### SLD microdissection

After a 6 week incubation period, mice were anesthetized using isoflurane (3% in 0.8L/minute of oxygen) until standard loss of reflexes, then rapidly decapitated (age: 12-14 weeks). The brain was removed from the skull and placed in ice cold calcium-free PBS for 3 minutes. After this, the brain was sectioned using a 1mm coronal brain matrix, and a slice containing the SLD was selected between −5.0 and −6.0mm posterior from bregma. Using a 0.75 mm tissue biopsy micropunch the SLD was microdissected and flash frozen on dry ice in a low-protein bind Eppendorf tube.

### Single nucleus RNA sequencing

SLD tissue samples (n=8 mice, male) were submitted to the 10X Genomics facility at the Princess Margaret Genomics Centre (Toronto, Canada). Nuclei isolation was performed by facility personnel, according to facility standards. Quality control was performed using a Luna Dual Fluorescence Cell counter to assess cell viability (<1%) and count nuclei. Libraries were created using the Chromium Single Cell 3’ Library Construction Kit v3 (10X Genomics, CAT# 1000190) according to the manufacturers protocol. After this cDNA was amplified for 11-13 PCR cycles and libraries were sequenced using the NextSeq 500 platform (Illumina) to a sequencing depth of ∼40,000 mean reads per nuclei.

### Custom transcriptome reference and data processing

FASTQ files were processed using the 10X Genomics CellRanger pipeline^58^ (version 8.0.0) and a custom transcriptome reference produced in house. The custom transcriptome reference was based on the GRCm38 Genome Assembly (mm10) with the nucleotide sequences for GFP (NIH GenBank Accession code OQ870305.1) and humanized ChR2 with H134R mutation fused to mCherry (SnapGene sequence #354478) were appended with the CellRanger mkref pipeline. Sparse count matrix files (.h5) were imported into R Studio for processing with Seurat^30^ v5^29^ using standard settings. Doublet removal was performed in Seurat with DoubletFinder^59^ and standard settings. Data filtration was performed for nFeature RNA (>500, <6500), nCount RNA (<30,000) and percent mitochondrial RNA (<3%). Seurat function SCTransform^60,61^ was used to normalize and scale raw count data and find variable features. PCA and UMAP were used for dimensionality reduction and visualization with the Louvain method for unsupervised clustering. The number of PCs and resolution were adjusted appropriately for each analysis and are summarized in Table 1. The Seurat function FindAllMarkers with standard settings (log2FC ≥ 0.25, minimum percent, 25%, Wilcoxon rank-sum test with the Bonferroni correction for adjusted p-value) was used to identify cluster-defining genes and perform differential gene expression analysis (*adjusted p-value < 0.05). All snRNA-seq analysis was performed with customized R scripts built in Seurat v5.

**Table 1.** PCA and resolution parameters used in snRNA-seq analyses.

| Dataset | nPCs | Resolution |
| --- | --- | --- |
| All nuclei | 21 | 0.5 |
| Snap25 neuronal nuclei* | 22 | 0.4 |
| Slc17a6 neuronal nuclei* | 21 | 0.4 |
| Gfp neuronal nuclei | 10 | 0.2 |
| mCherry neuronal nuclei | 10 | 0.3 |

\*after subsetting

### Nuclei subsetting

To subset neuronal nuclei, nuclei were with non-zero expression of *Snap25* or *Slc17a6* were selected from the dataset. Using PCA and the Louvain method, *Snap25* (**SFig. 1A**) or *Slc17a6* (**SFig. 1C**) nuclei underwent unsupervised clustering (*Snap25*: nPCs = 20, resolution = 0.2; *Slc17a6*: nPCs = 21, resolution = 0.4). The expression of cell type marker genes was overlaid onto identified clusters (**SFig. 1B, D**) and clusters with high expression of glial markers were removed. The subsetted populations were hence considered neuronal nuclei for downstream analysis (**Figs. 2, 3, 4**).

## Acknowledgments

We thank The Princess Marget Genomics Centre for their technical assistance with nuclei isolation. This work was supported by the Canadian Institutes of Health Research grant PJT-183876 (to J.P.), Natural Sciences and Engineering Research Council of Canada grant UTEA2025-HL (to J.P.), and National Institutes of Health grants R01NS118856 and R01NS073613 (to P.M.F.) and R01NS091126 (to E.A.).

