## Supplementary Figure 1 for "Molecular Logic of Cell Diversity and Circuit Connectivity in the REM Sleep Hub"

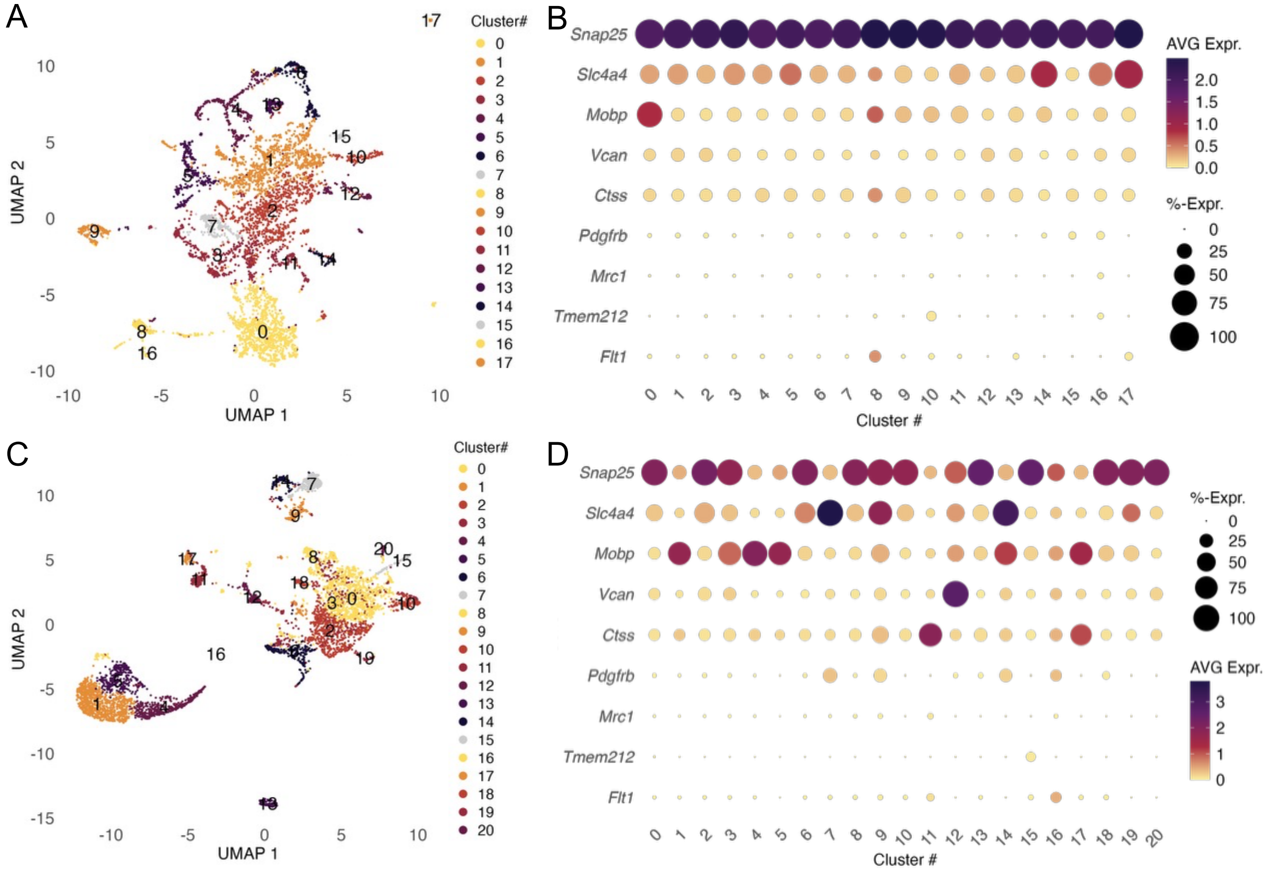


**Supplementary Figure 1. A.** UMAP plot for *Snap25*+ nuclei with PCA and unsupervised clustering. **B.** Dot plot depicting gene expression data for cell type marker genes, used to exclude glial nuclei and subset neuronal nuclei. **C.** UMAP plot for *Slc17a6*+ nuclei with unsupervised clustering. **D.** Dot plot depicting gene expression data for cell type marker genes, used to exclude glial nuclei and subset neuronal nuclei.
